# Identification and *In Silico* Characterization of Novel lncRNAs Associated with Dengue Infection

**DOI:** 10.1101/2025.05.08.652916

**Authors:** Abhay Deep Pandey, Jitendra kuldeep, Sudhanshu Vrati, Arup Banerjee

## Abstract

**Background:** Long noncoding RNAs (lncRNAs) are essential regulators of various biological processes in viral infections. In our previous study, we described the association of known lncRNAs with dengue disease progression. However, little is known about novel lncRNAs in the context of dengue infection. In this study, we aimed to identify novel lncRNAs and characterize their interaction with mRNAs during different stages of dengue infection using RNA sequencing (RNA-seq) technology to better understand their role in dengue pathogenesis.

**Methods:** We analyzed RNA-seq data (GSE94892) obtained from peripheral blood mononuclear cells (PBMCs) of 39 dengue-infected patients. RNA-seq data analysis was performed using the Tuxedo protocol, and the coding potential of novel transcripts was examined using CPC and CPAT tools. LncRNAs and cis-regulated mRNAs were identified through location-expression analysis within a 20 kbp genomic region surrounding the identified lncRNAs.

**Results:** We identified and characterized a total of 65 novel lncRNAs differentially expressed during dengue disease progression. These included 34 long intergenic lncRNAs, 2 antisense lncRNAs, and 25 alternatively spliced lncRNAs classified based on their localization. Bioinformatics analysis predicted potential cis-regulated target genes. Co-expression analysis revealed a strong correlation between lncRNAs and genes enriched in pathways related to megakaryocyte development and platelet production, suggesting a potential role in dengue pathogenesis. Furthermore, comparison with miRBase-21 indicated that some of the identified lncRNAs could act as miRNA precursors, highlighting a novel function of lncRNAs in gene regulation.

**Conclusion:** We developed an analytical pipeline to identify novel lncRNAs from RNA-seq data. Our analyses led to the identification of novel lncRNAs and their associated genes involved in dengue disease progression. These findings provide insights into the functional role of lncRNAs in dengue pathogenesis.

## Introduction

Dengue is the most prevalent mosquito-borne viral illness in humans, caused by the Dengue virus (DENV) [1]. While most dengue cases present as mild febrile illness, severe forms such as dengue hemorrhagic fever (DHF) and dengue shock syndrome (DSS) can lead to multiple organ failure and death, especially in children [2,3]. Dengue infection manifests in varying degrees of severity, ranging from mild febrile illness to severe disease with plasma leakage, shock, and multi-organ failure.

The pathogenesis of dengue virus infection is attributed to a complex interplay between the virus, host genes, and host immune response [19]. Host factors such as antibody-dependent enhancement (ADE), memory cross-reactive T cells, and genetic factors are major determinants of disease susceptibility. The pathophysiological hallmark of DHF/DSS is plasma leakage and deranged hemostasis, although the precise mechanism of this manifestation remains incompletely understood [20].

While protein-coding genes have been extensively studied, recent research has highlighted the importance of non-coding RNAs, which play regulatory roles in gene expression and impact various biological pathways, ultimately influencing disease outcomes [4]. Long noncoding RNAs (lncRNAs) are a class of RNAs longer than 200 nucleotides that do not encode proteins [5]. They are involved in numerous cellular processes, including gene regulation, post-transcriptional modifications, and epigenetic regulation. Recent studies have indicated that lncRNAs play a crucial role in viral infections, influencing host-virus interactions and immune responses [6].

LncRNAs are known to be involved in antiviral responses and various virus-host interactions, some of which may facilitate viral replication [7]. Cellular lncRNA expression can be altered in response to viral replication or viral protein expression. In our previous study, we identified several known lncRNAs induced during dengue virus infection [8]. One such lncRNA, NEAT1, was associated with dengue disease progression. Additionally, several novel transcripts were detected in PBMCs of dengue-infected patients. Here, we present an in-depth analysis of these novel lncRNAs, identifying their co-expression with mRNAs to reveal potential functional pathways impacted by their expression.

Recent studies have demonstrated that during DENV infection, heterogeneous nuclear ribonucleoproteins (hnRNPs) can regulate host mRNA expression, potentially influencing viral replication and pathogenesis [20]. The modulation of RNA-binding proteins by dengue infection can impact on mRNA transport and localization, which may be a critical factor in determining the course of the infection. Furthermore, the cytosolic localization of viral mRNA molecules can influence the accessibility of viral RNA to host immune factors, thereby affecting the recognition and clearance of the virus [20].

## Materials and Methods

### Patient Information and RNA Sequencing Data

RNA-seq data from peripheral blood mononuclear cells (PBMCs) of 39 dengue-infected patients were obtained from the Gene Expression Omnibus (GEO) database (accession number: GSE94892). Patients were classified based on clinical parameters according to the World Health Organization (WHO) guidelines into three groups: dengue without warning signs (DWS), dengue with warning signs (DWWS), and severe dengue (SD). Detailed patient demographics, clinical characteristics, and laboratory findings have been previously reported [9,10,11].

Total RNA was extracted from PBMCs using the RNeasy Mini Kit (Qiagen, Hilden, Germany) following the manufacturer’s protocol. RNA integrity was assessed using the Agilent 2100 Bioanalyzer (Agilent Technologies, Santa Clara, CA, USA), and samples with RNA integrity number (RIN) values >7.0 were selected for sequencing. RNA-seq libraries were prepared without poly-A selection to enable the detection of both polyadenylated and non-polyadenylated transcripts. Sequencing was performed on the Illumina HiSeq 2500 platform (Illumina, San Diego, CA, USA) with 2 × 100 bp paired-end reads, generating approximately 30-40 million reads per sample.

### RNA-seq Data Analysis and Identification of Novel Transcripts

Raw sequencing reads were quality-checked using FastQC (v0.11.9) and trimmed using Trimmomatic (v0.39) to remove adapters and low-quality bases (Phred score <20). Clean reads were aligned to the human reference genome (GRCh38/hg38) using HISAT2 (v2.2.1) with default parameters. The resulting alignment files were processed using StringTie (v2.1.4) to assemble transcripts, followed by Cufflinks (v2.2.1) to merge assemblies across all samples.

The assembled transcripts were compared with the reference annotation (GENCODE v38) using Cuffcompare to identify novel transcripts. Transcripts with class codes ‘i’ (intronic), ‘u’ (intergenic), and ‘x’ (antisense) were retained for further analysis, while those annotated as known genes (‘=‘) were excluded [12,13,14]. To ensure high-quality novel transcripts, we applied the following filtering criteria:

1. Transcript length ≥200 nucleotides
2. Number of exons ≥2
3. Expression level ≥1 FPKM (fragments per kilobase of transcript per million mapped reads) in at least one sample
4. Maximum open reading frame (ORF) length <120 amino acids (360 nucleotides)

ORF length was evaluated using TransDecoder (v5.5.0), and transcripts with ORFs longer than 120 amino acids were excluded to minimize the inclusion of potential protein-coding genes.

### Coding Potential Assessment

To ensure that the identified novel transcripts were non-coding, we assessed their coding potential using two complementary tools: Coding Potential Calculator (CPC) [13] and Coding-Potential Assessment Tool (CPAT) [14]. CPC evaluates the coding potential based on sequence features and the quality of the predicted ORF, while CPAT uses a logistic regression model based on sequence features such as ORF length, ORF coverage, Fickett score, and hexamer usage bias.

For CPC analysis, we used the web-based CPC2 server with default parameters. Transcripts with a CPC score <0 were classified as non-coding. For CPAT analysis, we used the human-specific model with a cutoff value of 0.364, below which transcripts were classified as non-coding. Only transcripts classified as non-coding by both tools were retained for subsequent analyses.

### Differential Expression Analysis

Differential expression analysis of known genes, novel genes, and lncRNAs was performed using Cuffdiff (v2.2.1). The expression levels were calculated in FPKM, and differentially expressed (DE) genes and lncRNAs were identified based on an adjusted p-value <0.05 (Benjamini-Hochberg correction for multiple testing) [16]. Hierarchical clustering of DE lncRNAs and genes was performed using the hclust function in R (v4.1.0) with Euclidean distance and complete linkage method to visualize expression patterns among samples. The results were visualized using the pheatmap package in R.

### Functional Annotation and Pathway Enrichment Analysis

To infer the potential functions of novel lncRNAs, we performed co-expression analysis and cis-regulation analysis. For co-expression analysis, Pearson and Spearman correlation coefficients were calculated between each lncRNA and all protein-coding genes across all samples. Gene pairs with correlation coefficients >0.9 and p-values <0.05 were considered significantly co-expressed.

For cis-regulation analysis, we identified protein-coding genes located within a 20 kbp window upstream and downstream of each lncRNA [18]. The expression correlation between lncRNAs and their neighboring genes was calculated as described above, and gene pairs with significant correlations were considered potential cis-regulatory relationships. This approach is based on the understanding that lncRNAs can act as spatial amplifiers that control nuclear structure and gene expression, particularly in regulating nearby genes [17].

Gene Ontology (GO) and Kyoto Encyclopedia of Genes and Genomes (KEGG) pathway enrichment analyses were performed using the clusterProfiler package in R for the co-expressed and cis-regulated genes. Enriched GO terms and KEGG pathways with adjusted p-values <0.05 were considered significant. The results were visualized using the enrichplot package in R and Cytoscape (v3.6.0) for network visualization.

### Evolutionary Conservation Analysis

To assess the evolutionary conservation of the identified lncRNAs, we used the phastCons and phyloP scores from the UCSC Genome Browser, which measure sequence conservation across multiple vertebrate species. For each lncRNA, we calculated the average phastCons and phyloP scores across the entire transcript length and compared them with those of protein-coding genes and random genomic regions.

LncRNAs with high conservation scores were further analyzed for potential functional domains using HMMER against the Pfam database [15].

### Secondary Structure Prediction and Functional Motif Identification

RNA secondary structure plays a crucial role in lncRNA function. We used the RNAfold algorithm from the ViennaRNA package to predict the secondary structures of the identified lncRNAs. Additionally, we searched for known RNA binding protein (RBP) motifs using the RBPmap web server and RBP interaction data from ENCODE, focusing on RBPs known to be involved in immune response and viral infection.

### Genomic Distribution Analysis

To visualize the chromosomal distribution of the identified lncRNAs, we generated Circos plots mapping each lncRNA to its genomic location. This analysis allowed us to identify potential clustering patterns and chromosomal enrichment of dengue-associated lncRNAs.

## Results and Discussion

### Identification of Novel lncRNAs in Dengue-Infected Patients

Using our computational pipeline (Figure 1A), we identified a total of 65 novel lncRNAs that were differentially expressed during dengue disease progression. These lncRNAs were classified based on their genomic context: 34 long intergenic lncRNAs (lincRNAs) located between protein-coding genes, 2 were antisense lncRNAs transcribed from the opposite strand of protein-coding genes, and 25 were alternatively spliced lncRNAs generated through alternative splicing events.

**Figure 1.**
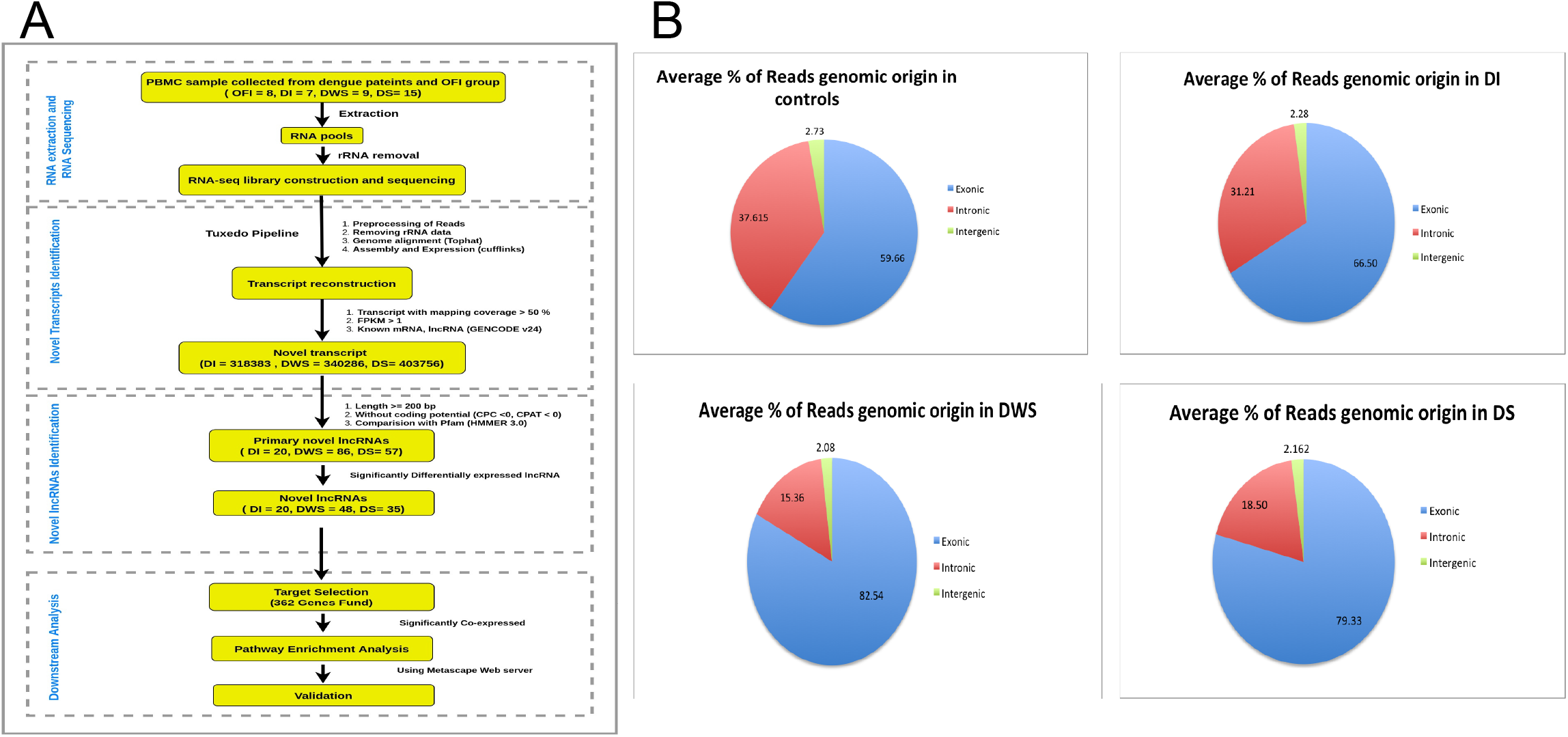
Identification of novel lncRNAs in dengue-infected patients. (A) Computational pipeline for the identification of novel lncRNAs from RNA-seq data. The workflow includes RNA extraction, transcript reconstruction, novel transcript identification, and lncRNA validation. (B) Genomic origin distribution of RNA-seq reads in different patient groups: Other Febrile Illness (OFI), Dengue Infection (DI), Dengue with Warning Signs (DWS), and Dengue Severe (DS). The pie charts show the percentage of reads mapping to exonic, intronic, and intergenic regions.

Analysis of the genomic origin of RNA-seq reads revealed distinct patterns across different patient groups (Figure 1B). In the Other Febrile Illness (OFI) control group, 59.66% of reads mapped to exonic regions, 37.61% to intronic regions, and 2.73% to intergenic regions. In contrast, dengue infection groups showed a progressive increase in exonic mapping: 66.50% in Dengue Infection (DI), 82.54% in Dengue with Warning Signs (DWS), and 79.33% in Dengue Severe (DS). This shift in genomic origin of transcription suggests widespread changes in gene expression and alternative splicing during dengue infection progression.

### Differential Expression Patterns of Novel lncRNAs

The expression profiles of these novel lncRNAs showed distinct patterns across different dengue severity groups (Figure 2A). Hierarchical clustering analysis revealed three main clusters of lncRNAs: (1) those upregulated in severe dengue, (2) those upregulated in dengue with warning signs, and (3) those downregulated in severe dengue. These distinct expression patterns suggest that different sets of lncRNAs may play specific roles in different stages of dengue disease progression.

**Figure 2.**
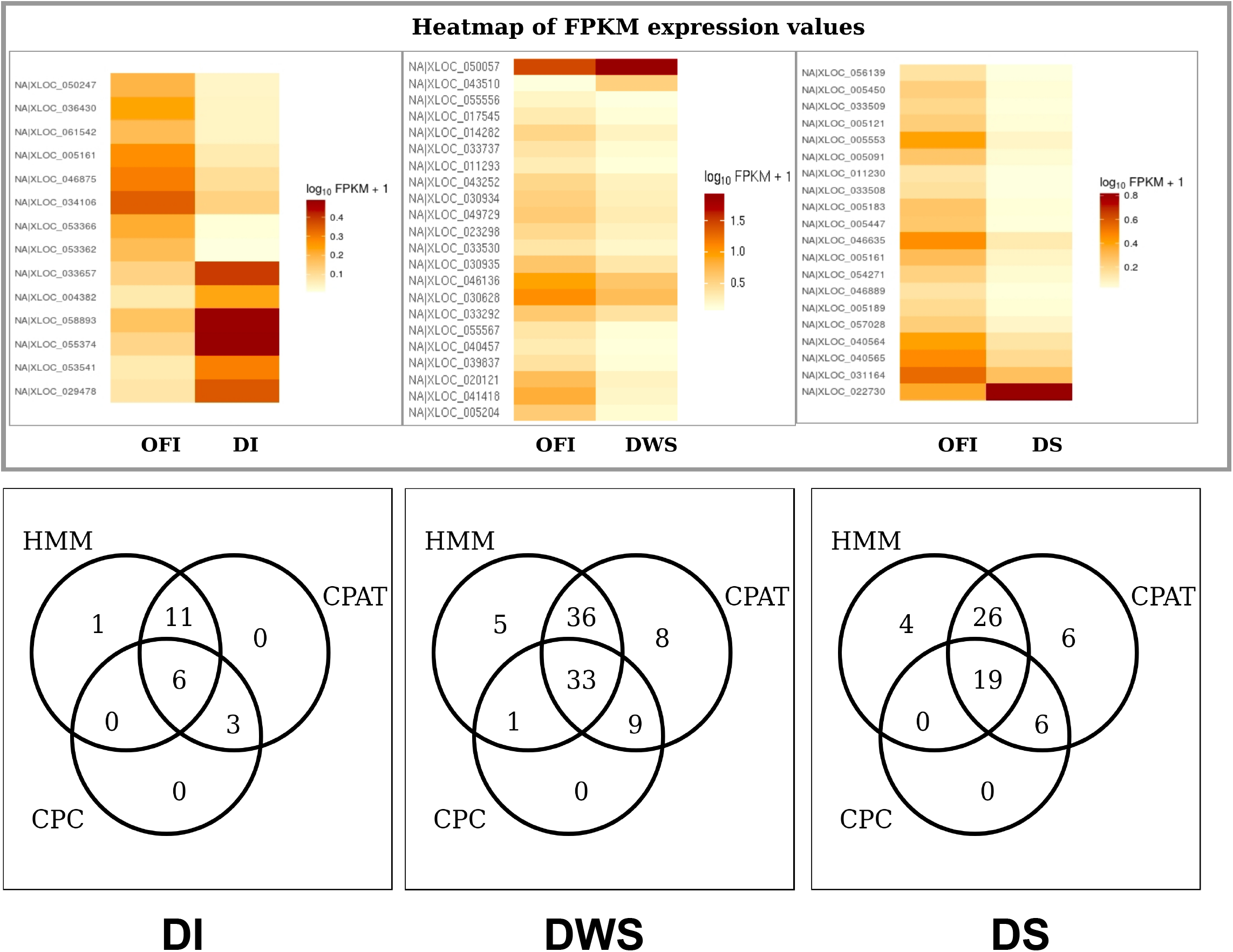
Characterization and validation of novel lncRNAs. (A) Heatmap showing differential expression of novel lncRNAs across dengue severity groups (OFI vs. DI, OFI vs. DWS, OFI vs. DS). Color intensity represents log10-transformed FPKM values, with darker colors indicating higher expression. (B) Venn diagrams illustrate the consensus between three computational methods (HMM, CPAT, and CPC) for assessing non-coding potential of transcripts identified in DI, DWS, and DS patients.

To validate the non-coding nature of the identified transcripts, we employed multiple computational tools (Figure 2B). The Venn diagrams show the overlap between predictions from three methods: HMM (HMMER), CPAT, and CPC. In the DI group, 6 transcripts were identified as non-coding by all three methods. In the DWS group, 33 transcripts were predicted as non-coding by all methods, while 19 transcripts in the DS group were classified as non-coding by all tools. This consensus approach ensured high confidence in our lncRNA identification.

### Functional Characterization of Novel lncRNAs

To investigate the potential functions of the identified novel lncRNAs, we performed co-expression analysis with protein-coding genes. We constructed co-expression networks for each dengue severity group (Figure 3A). In the DI group, several lncRNAs showed strong co-expression with chemokine receptors CXCR1 and CXCR2, as well as ALSCR11, suggesting potential roles in immune response and cell migration. The DWS network revealed interactions with genes involved in transcriptional regulation (EGR1), immune signaling, and cellular stress response. In the DS network, co-expression with genes related to inflammatory response (C1GA14), transcription regulation (TCF3, ZBP1), and cellular transport were observed, highlighting the complex regulatory roles of these lncRNAs in severe dengue.

**Figure 3.**
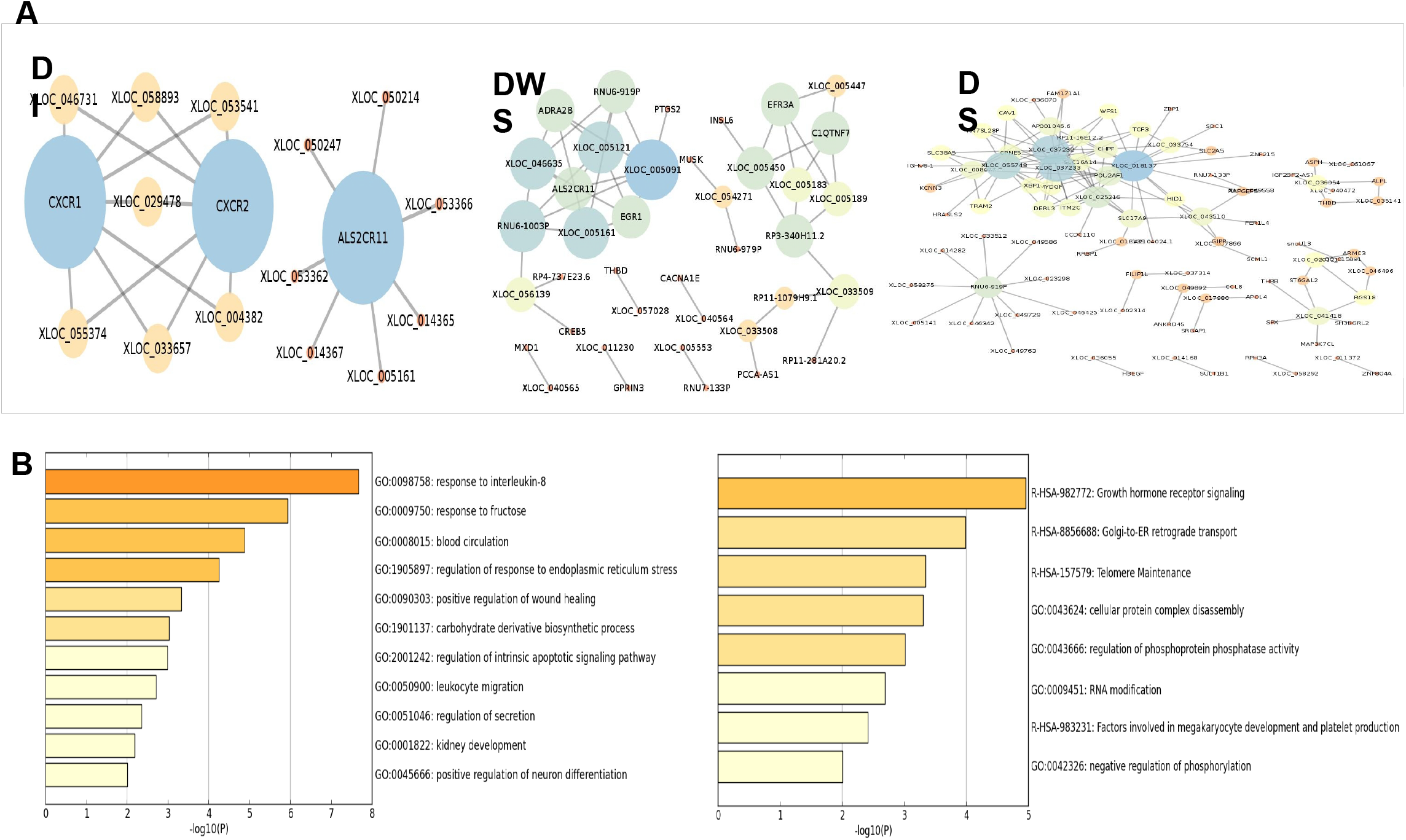
Co-expression network and pathway analysis of novel lncRNAs. (A) Co-expression networks showing interactions between novel lncRNAs (yellow nodes) and protein-coding genes (blue/green nodes) in DI, DWS, and DS patients. Edge connections represent significant correlation in expression patterns. (B) Enriched biological processes (top) and pathways (bottom) associated with genes co-expressed with novel lncRNAs. Bar length represents statistical significance (−log10(P-value)).

Functional enrichment analysis of co-expressed genes revealed significant associations with biological processes and pathways related to immune response, platelet function, and vascular permeability (Figure 3B). GO analysis identified enrichment in response to interleukin-8, blood circulation, regulation of endoplasmic reticulum stress, positive regulation of wound healing, and leukocyte migration, among others. These processes are highly relevant to dengue pathogenesis. Pathway analysis revealed enrichment in growth hormone receptor signaling, Golgi-to-ER retrograde transport, telomere maintenance, and notably, megakaryocyte development and platelet production, which is particularly significant given that thrombocytopenia is a hallmark of severe dengue.

The genomic distribution analysis revealed in our Circos plots (Figure 4A-C) provides additional context for understanding the potential regulatory relationships between these lncRNAs and their neighboring genes. Many of the identified lncRNAs are in close proximity to immune-related genes, suggesting possible cis-regulatory mechanisms. For example, several lncRNAs on chromosome 6 map near the major histocompatibility complex (MHC) region, which is critical for immune response. Similarly, the cluster of lncRNAs on chromosome 13 is located near genes involved in platelet function, supporting our co-expression analysis findings.

**Figure 4.**
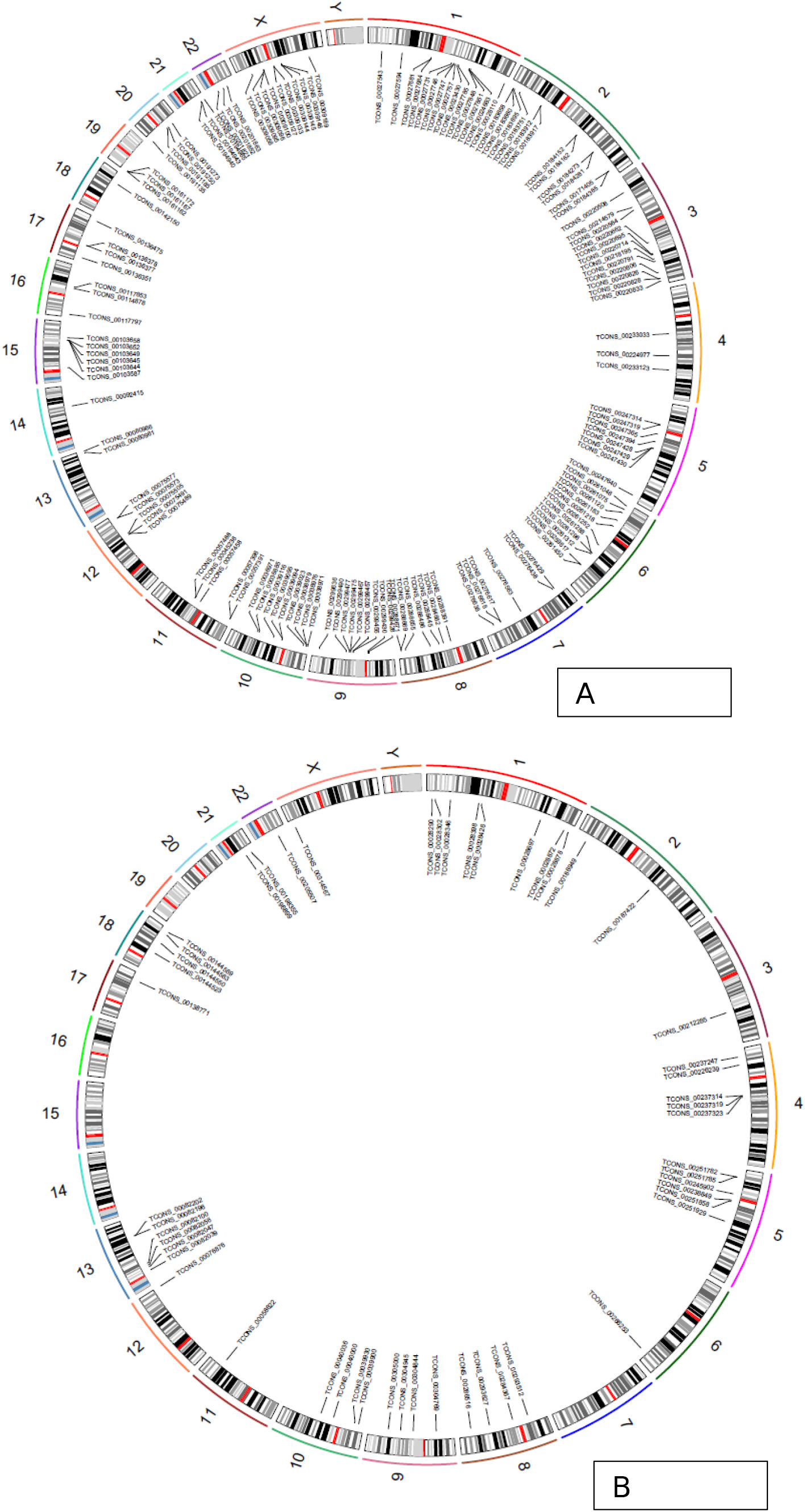
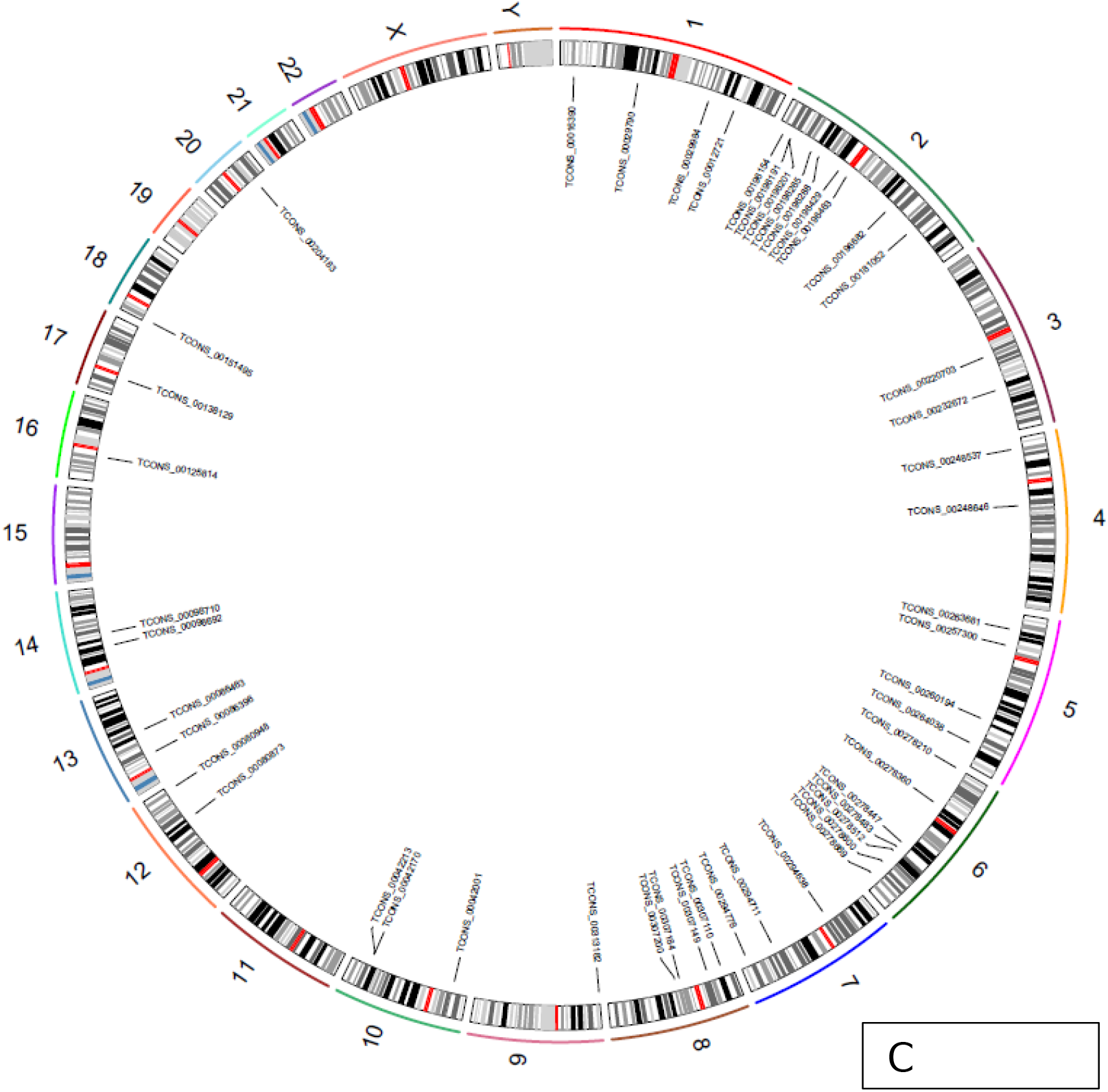
Genomic distribution of novel lncRNAs associated with dengue infection. (A) Circos plot showing the chromosomal locations of all 65 differentially expressed lncRNAs across the human genome. (B) Circos plot illustrating the distribution of lncRNAs with the strongest differential expression between dengue severity groups. (C) Circos plot depicting the genomic locations of lncRNAs significantly co-expressed with immune-related genes and platelet production pathways. Chromosomal ideograms are shown in the outer ring, with lncRNA transcripts indicated by their unique identifiers.

### Novel lncRNAs as Potential MicroRNA Precursors

Comparison of the identified lncRNAs with miRBase-21 revealed that 12 of the 65 novel lncRNAs showed significant sequence similarity to known miRNA precursors. Secondary structure analysis confirmed that these lncRNAs could form stable hairpin structures typical of miRNA precursors, suggesting a potential role as miRNA precursors.

This finding highlights a novel function of lncRNAs in gene regulation during dengue infection, as they may not only directly regulate gene expression through cis- or trans-acting mechanisms but also serve as precursors for miRNAs, which in turn can regulate multiple target genes post-transcriptionally.

### Evolutionary Conservation and Secondary Structure Analysis

Analysis of evolutionary conservation showed that most of the identified lncRNAs exhibited low sequence conservation across vertebrates, consistent with the general observation that lncRNAs are less conserved than protein-coding genes. However, several lncRNAs showed highly conserved regions, suggesting potential functional importance. Interestingly, the lncRNAs with the highest conservation scores were also those most strongly associated with immune response pathways in our co-expression analysis.

Secondary structure prediction revealed that many of the identified lncRNAs contained stable stem-loop structures, which could serve as binding sites for RNA-binding proteins or other regulatory molecules. Analysis of known RBP motifs identified potential binding sites for several immune-related RBPs, including hnRNPs, which have been shown to be modulated during dengue infection [20]. This finding suggests that these lncRNAs may function by interacting with RBPs to regulate immune response genes during dengue infection.

### Role of lncRNAs in Dengue Virus-Host Interactions

Understanding the complex interactions between DENV and host molecular machinery is essential for developing new therapeutic strategies. Recent studies have shown that DENV can modulate host epigenetic regulation through interactions with various host cellular genes and lncRNAs, resulting in RNA methylation and other post-transcriptional modifications [21]. Our findings contribute to this growing body of evidence by identifying novel lncRNAs potentially involved in these regulatory processes.

The cross-talk between viral and host factors during DENV infection is a critical determinant of disease outcome. Research has demonstrated that DENV can inhibit type I interferon production through its NS2B-NS3 protease complex, which targets adaptor molecules for degradation [23]. Our co-expression analysis identified several lncRNAs significantly correlated with genes involved in interferon signaling pathways, suggesting a potential role for these lncRNAs in modulating the host antiviral response.

Furthermore, recent findings have highlighted the importance of subgenomic flaviviral RNA (sfRNA) in dengue virus replication and host immune evasion [24]. SfRNA is derived from the 3’ untranslated region (UTR) of the DENV genome and can interact with host proteins to suppress antiviral immune activation. Interestingly, several of our identified lncRNAs shared sequence similarity with regions of the DENV 3’ UTR, suggesting potential competitive or cooperative interactions with sfRNA in regulating host gene expression.

The Circos plots we generated (Figure 4A-C) reveal that the genomic distribution of dengue-associated lncRNAs is not random but shows distinct patterns of enrichment on specific chromosomes. This spatial organization may reflect functional significance in the host response to DENV infection. The chromosomal regions with high densities of dengue-responsive lncRNAs may represent genomic hotspots for host-virus interactions, potentially serving as important regulatory hubs during infection.

### Comparison with lncRNAs in Other Vector-Borne Diseases

To place our findings in a broader context, we compared the identified lncRNAs with those reported in other vector-borne diseases. A recent study identified 2,949 lncRNAs in the malaria mosquito vector, Anopheles gambiae, with considerably lower sequence conservation compared to protein-coding genes [24]. Similarly, another study reported that 43% of total midgut transcripts of An. gambiae are lncRNAs, with 32% showing some level of homology to other species [26].

In Aedes aegypti, the primary vector for dengue virus, a comprehensive list of long intergenic non-coding RNAs has been reported, with some differentially expressed in mosquitoes infected with dengue virus [26]. These lncRNAs could be involved in DENV-mosquito interactions and provide new avenues for exploring vector biology and virus-vector interactions. Our study extends these findings to human host cells, providing insights into the role of lncRNAs in DENV-human interactions.

### Implications for Dengue Pathogenesis and Potential Therapeutic Targets

The pathogenesis of dengue virus infection is complex and involves multiple factors, including viral replication, immune response, platelet dysfunction, and vascular leakage. Our findings suggest that lncRNAs may play important roles in these processes, particularly in immune regulation and platelet production.

Thrombocytopenia and platelet dysfunction are commonly observed in DENV patients, and dengue complications are usually preceded by a rapid drop in platelet count [28]. Platelets play key roles not only in coagulation but also in inflammation, host defense, and preservation of endothelial integrity, especially under inflammatory conditions. Our co-expression analysis identified several lncRNAs strongly correlated with genes involved in platelet production and function, suggesting a potential role in dengue-associated thrombocytopenia.

Furthermore, endothelial activation and increased vascular permeability are hallmarks of severe dengue. Recent studies have shown that von Willebrand factor (VWF) levels are elevated in dengue infection, particularly in DSS patients [28]. VWF is released from endothelial cells upon activation and plays a crucial role in platelet adhesion and aggregation. Our analysis identified lncRNAs co-expressed with genes involved in endothelial cell function and vascular permeability, suggesting potential targets for therapeutic intervention.

In the absence of specific antiviral drugs for dengue, targeting host factors required for viral replication has emerged as an alternative strategy [23]. LncRNAs that modulate host-virus interactions could serve as potential therapeutic targets. For instance, silencing of lncRNAs that facilitate viral replication or enhance immune dysfunction could potentially mitigate disease severity.

The genomic distribution analysis presented in our Circos plots (Figure 4A-C) provides valuable insights into identifying potential therapeutic targets. By highlighting chromosomal regions enriched with dengue-responsive lncRNAs, these visualizations may guide future investigations into key regulatory networks involved in dengue pathogenesis. The lncRNAs clustered in immune-related genomic regions, as shown in Figure 4C, represent particularly promising candidates for therapeutic intervention.

## Conclusion

In this study, we developed a computational pipeline to identify and characterize novel lncRNAs from RNA-seq data of dengue-infected patients. We identified 65 novel lncRNAs differentially expressed during dengue disease progression and provided insights into their potential functions through co-expression and cis-regulation analyses.

Our findings suggest that these novel lncRNAs may play important roles in dengue pathogenesis by regulating genes involved in immune response, platelet production, and vascular permeability. Furthermore, some of these lncRNAs may function as miRNA precursors, adding another layer of complexity to gene regulation during dengue infection.

The genomic distribution analysis revealed distinct patterns of lncRNA localization across the human genome, with notable enrichment on specific chromosomes. These spatial patterns provide additional context for understanding the regulatory relationships between lncRNAs and their target genes during dengue infection.

This study expands our understanding of the lncRNA landscape in dengue infection and provides a foundation for future functional studies to elucidate the precise mechanisms by which these lncRNAs contribute to dengue pathogenesis. Such knowledge could lead to the development of novel diagnostic markers or therapeutic targets for dengue, a disease with significant global health impact.

The complex interplay between lncRNAs, viral factors, and host immune response represents a new frontier in understanding dengue pathogenesis. Future studies should focus on validating the functional roles of these lncRNAs *in vitro* and *in vivo*, as well as exploring their potential as biomarkers for disease severity and progression. Additionally, investigating the conservation and function of these lncRNAs across different dengue serotypes and in related flavivirus infections could provide broader insights into the role of non-coding RNAs in arboviral diseases.

